# A first morphological and electrophysiological characterization of Fañanas cells of the mouse cerebellum

**DOI:** 10.1101/2023.09.26.559486

**Authors:** A. Singer, L. Vinel, F. Trigo, I. Llano, M. Oheim

**Author notes:** **Correspondence** Martin Oheim, Université Paris Cité, CNRS, Saints-Pères Paris Institute for the Neurosciences, F-75006 Paris, France.

## Abstract

The “feathered” cells of Fañanas (FCs) are cerebellar glia of unknown function. Initially described more than a century ago, they have been practically absent from the scientific literature. Here, we combined whole-cell patch-clamp recordings, dye loading and near UV-laser photolysis for a first characterization of FCs in terms of their morphology, electrophysiological properties and glutamate-evoked currents. FCs were identified in cerebellar slices by their small cell bodies located in the molecular layer and stubby processes. Despite a more compact shape compared to Bergmann glia (BGs) FCs had similar membrane resistances and basal currents, suggesting electrical coupling between neighboring glia. Dye filling and pharmacological experiments confirmed homo- and heterotypic gap junction coupling among FCs and BGs. Parallel-fiber stimulation evoked in FCs a slow inward current partially blocked by NBQX, along with superimposed fast (ms) transients. Repetitive stimulations resulted in a rapid desensitization of this AMPA-receptor mediated current, which recovered for stimulation intervals >0.5s. Laser photolysis of MNI-caged glutamate replicated fast inward currents with highest current densities in the distant process. We conclude that FCs respond with fast AMPA currents to local glutamate release, and that they integrate ambient glutamate to a slow inward current. Interestingly, FCs prevail throughout adulthood with markedly different densities among cerebellar lobes. Thus, FCs are not just displaced BGs as previously suggested, but they may have lobule specific functions, both locally and at the circuit level, yet to be uncovered.

## 1 INTRODUCTION

The cerebellar circuitry is one of the best characterized of the entire brain, both from an anatomical and functional standpoint. Strangely, this is true with one exception: the feathered cell of Fañanas, a cerebellar astrocyte discovered over a century ago [Fañanas (1916)], but that has been practically absent from the scientific literature since.

Cerebellar glia were initially described by Bergmann (1857), Golgi (1885), Gierke (1886), Cajal (1911) as well as Jorge Ramón y Fañanas(1916), among others. Velate astrocytes can be divided into Golgi epithelial cells and protoplasmic astrocytes of the granule cell layer. The first type gives rise, during development, to the Bergmann fibers that make contact with PCs. The second type forms a sheath around granule cells, dividing the granular layer into distinct compartments containing individual glomeruli and single or clustered granule cells [Chan-Palay and Palay (1972)]. Most research on cerebellar glia has focused on Bergmann glia (BGs), while velate astrocytes of the granule-cell layer have received comparatively less attention [Hoogland and Kuhn (2010); Chan-Palay and Palay (1972)]. BGs are specialized unipolar astrocytes that exhibit close developmental, anatomical, and functional relationships with Purkinje cells (PCs) [Bellamy (2006), Hoogland and Kuhn (2010)]. The candelabrum-shaped processes of BGs co-align with PC dendrites, providing a structural scaffold for their directed growth during development [Lordkipanidze and Dunaevsky (2005)]. In contrast to most other rodent astrocytes, the arborizations of BGs interdigitate in the adult brain [Grosche et al. (2002)]. Electrical stimulation of parallel fibers (PFs) induces complex calcium (Ca^2+^) signals in BGs, ranging from localized Ca^2+^ transients in subcellular microdomains [Grosche et al. (1999)] to large-scale Ca^2+^ waves that invade extensive networks of BGs. *In vivo*, BGs exhibit synchronous activation, e.g. during head-turning and activity of BGs elicits measurable changes in brain dynamics or blood flow [Nimmerjahn et al. (2009)].

A largely forgotten type of cerebellar glia needs to be added to this picture, the “feathered” cell of Fañanas (FC), a compact, unipolar astrocyte with a thick process decorated with numerous fine and short protrusions [Fañanas (1916); Goertzen and Veh (2018)]. FCs have been visualised originally through a tedious goldsublimate procedure [Fañanas (1916)]. The origin, development and function of FCs in the adult cerebellum are unknown. Recent work [Goertzen and Veh (2018)] identified, in rat, two potassium channel-related polypeptides, Kv2.2 and Calsenilin (KChIP3), respectively, as potential FC-specific marker proteins. We reasoned that their specific expression in FCs should allow us addressing optogenetic tools for reading out or stimulating selectively FCs activity in a transgenic mouse model, and thus pave the way for the functional studies of FCs with modern techniques. However, while we could replicate Goertzen’s and Veh’s findings in rat, identical experiments in mouse failed to produce a specific labelling. These early experiments allowed us to familiarise ourselves with FCs of the molecular layer to a degree that we could routinely identify them on differential interference contrast (DIC) images.

In the present study, we used whole-cell patchclamp recordings, extracellular field stimulations, near-UV-glutamate uncaging, imaging of dye-coupling, and morphometry of reconstructed 3-D confocal images to provide the first physiological characterization of Fañanas glia. In Aldh1L1-GFP transgenic mice, we detected FCs from P6 and throughout adulthood. We find that FCs share many functional characteristics with neighboring BGs, including their passive membrane properties, a resting potential close to the potassium reversal potential and functional AMPA-receptor expression. Albeit not rare, FCs are less abundant than BGs and clearly morphologically distinct. The ability of FCs to read out millisecond glutamate release events as well as their gap-junction coupling to neighboring BGs and FCs suggest specific, yet to uncover, functions in cerebellar signal processing. In another paper (Singer*, Vinel*, et al, *in preparation*), we will provide a comprehensive molecular characterization of FCs using immunofluorescence combined with confocal and STED imaging, and we also expand our analysis to the FC population intermingled in the PC layer.

## 2 MATERIAL AND METHODS

Additional experimental procedures are found in the Supporting Information (SI), online.

### 2.1 Drugs and pharmacological agents

Most chemical products including Carbenoxolone (7421-40-1), NBQX (118876-58-7) and Spermine (7144-3) were purchased from Sigma-Aldrich (Saint-Louis, US). MNI-glutamate (HB0423) was from HelloBio (Bristol, UK), and Alexa-594 (A10438) from Thermo Fisher Scientific (Waltham, US).

### 2.2 Animals

All experiments followed European Union (EU) and institutional (CNRS or IIBCE, respectively) guidelines for the care and use of laboratory animals (Council Directive 2010/63/EU and CEUA approval number 001-01-2023, respectively). This study used a total of 10 adult (postnatal day P30-P120) C57BL/6J mice (Paris) and 25 BALB/c mice (P25-P35, Montevideo), as well as 100 transgenic Aldh1L1-GFP mice [Cahoy et al. (2008)], aged P6 to P200. Mice were housed in an enriched environment (Paris) at constant cage temperature (22°C) with *ad libitum* access to food and social interactions. For reproducing the results from [Goertzen and Veh (2018)], AS worked in Rüdiger Veh’s lab at the Charité (Berlin, Germany), where we used two rats. No distinction was made between male or female rodents.

### 2.3 ALDH1-EGFP fluorescence and immunochemistry

#### 2.3.1 Brain removal

Mice were deeply anesthetized using ketamine-xylazine via intraperitoneal (i.p.) injection in accordance to their weight. Anesthetized animals were positioned in a *dorsal decubitus* position on the operating table, with their feet pinned. A thoracic flap was created and reclined to expose the heart. Subsequently, a needle was inserted into the apex of the left ventricle, and the right atrium was incised to allow the circulation to flow out of the body. The blood circuit was then washed by perfusing phosphate-buffered saline (PBS) (150 mM NaCl in 10 mM phosphate buffer, pH 7.4). Care was taken to avoid bubble formation. PBS was replaced with a solution of 4% paraformaldehyde (PFA) in PBS. The perfusion was considered complete when the tail or hind paw of the mouse became rigid. The brains were removed and placed in a 4% PFA solution at 4 °C for 3 hours.

#### 2.3.2 Slice preparation

After several washes with PBS, the cerebellar vermis was extracted and glued to the holder of a vibratome (Leica VT1000S, Wetzlar, Germany). 200-μm sagittal slices were cut in PBS and stored in 1x PBS at 4 °C.

#### 2.3.3 Tissue clearing

For imaging fine tissue detail we used chemical CUBIC tissue clearing [Matsumoto et al. (2019)]. After two 30-min washes with PBS 1x, the slices were incubated during 2h in CUBIC-L solution, and thereafter thoroughly washed three times during 20 min each with PBS. Slices were mounted between slides and coverslips using spacers, and the chamber volume was filled up with 10 μL of CUBIC-R solution.

#### 2.3.4 Confocal imaging

Three-dimensional (3-D) fluorescence images were acquired on a Zeiss LSM710 META confocal microscope, using either a x20/NA0.8 air objective (working distance, WD: 0.55 mm) or a x63/NA1.4 oil immersion lens (WD: 0.19 mm), respectively. We employed ZEN Black 2 software for acquisition and pre-processing. The confocal pinhole diameter was systematically set to 1 Airy unit. In multicolor acquisitions, images were acquired sequentially using single-wavelength excitation to prevent cross-excitation and bleed-through.

### 2.4 Acute slice preparation for electrophysiology and dye-filling experiments

We prepared 200-μm thick sagittal slices from cerebellar cortices on a vibratome (either a VT1200S, Leica, Wetzlar, Germany, or a Microm HM650V Thermo Fisher Scientific, Waltham, US) using published procedures [Alcami et al. (2012)]. Compositions of the cutting, recovery and recording solutions were identical, each containing (in mM): 130 NaCl, 2.5 KCl, 1.3 NaH_2_PO_4_, 26 NaHCO_3_, 10 glucose, 2 CaCl_2_, 1 MgCl_2_. Osmolarity was 306 mOsm/l) and solutions were bubbled with a 95% O_2_ / 5% CO_2_ mix to have a pH of 7.4. Slices were prepared in this solution at 4 °C and kept thereafter at 34 °C to recover. Pipettes were pulled from borosilicate glass with filament (BF150-86-7.5HP, Sutter Instrument, Novato, US) on a vertical puller (either a PC-100, Narashige, Tokyo, Japan, or a PIP-6, HEKA Elektronik, Stuttgart, Germany) and had 7.5-8.5 MΩ resistance when filled with K-gluconate solution. We did not polish pipettes after pulling. For whole-cell patch-clamp recordings, we used standard upright IR-DIC microscopes (either a Zeiss AXIO OBSERVER, Oberkochen, Germany, or an Olympus BX51, Tokyo, Japon), fitted with water immersion PlanApo ×63/NA0.9w objective or water immersion ×60/NA1.0w objective and a CCD camera (Cohu, Poway, CA). Currents were recorded and digitized with an EPC-9 patch clamp amplifier (HEKA Multi Channel Systems MCS GmbH, Reutlingen, Germany) controlled with PatchMaster software. We used 3-axes micromanipulators (Luigs and Neumann GmbH, Ratingen, Germany) for positioning the patch and stimulation electrodes. Bath solutions were exchanged by aid of Peristaltic perfusion system using a PPS2 pump (MCS, Reutlingen, Germany). Unless otherwise stated, all experiments were performed at room temperature (RT, 22-23°C). In some experiments, we used a Peltier stage (EA Elektro-Automatik, PS 2016-100, Viersen, Germany) to maintain the bath temperature at 34 °C. Intracellular (pipette) solution contained (in mM): 144 K-gluconate, 6 KCl, 4.6 MgCl_2_, 10 HEPES, 1 EGTA, 0.1 CaCl_2_, 4 NaATP, 0.4 Na-GTP. For voltage-clamp recordings, BGs and FCs were held at -90 mV. I-V curves were recorded by applying 20-ms pulses to values ranging from -100 mV to 40 or 0 mV. Potentials were corrected for -12 mV liquid junction potential. Where indicated, we added to the pipette solution 20 μl of Alexa-594 (ex./em. = 588 nm/613 nm; Thermo Fisher Scientific, Waltham, US) for the morphological reconstruction of the FCs previously recorded from. The pipette was withdrawn, the slice immediately fixed with 4% PFA, and the 3-D morphology reconstructed from serial optical sections of confocal micrographs using the ×63/NA1.0w objective. Where indicated, we added 2 μM NBQX to block AMPA receptors, 0.8 mM spermine to open gap junctions [Skatchkov et al. (2015); Benedikt et al. (2012)] or 150 μM carbenoxolone (CBX) as a gap-junction blocker [Davidson et al. (1986)].

### 2.5 Photolysis of caged MNI-glutamate

Glutamate was photolysed from 4-Methoxy-7-nitroindoli-inyl-caged-L-glutamate (MNI-GLutamate, HelloBio, Bristol, UK) by focusing the beam of a 405-nm diode laser (Obis, Coherent, California, USA) through a ×60/NA1.0w objective (Olympus, Hamburg, Germany) to a 1 μm diameter spot in the sample plane (Fig. S1) [Trigo et al. (2009); Zorrilla de San Martin et al. (2017)]. MNI-Glutamate was added to the bath at a final concentration of 0.5 to 1 mM and released with 100- to 500-μs light pulses at 1- to 3-mW laser power. We did not note any noxious side-effects from photolysis by-products. The power of the laser-pulse was monitored on a calibrated photodiode. In our setup, the laser spot is fixed, and the photolysis location is chosen by positioning the slice on the region of interest (ROI). Before each experiment, the exact location of the laser spot was verified and the cell’s position corrected if necessary.

### 2.6 Data analysis and statistics

We used IGOR8 (Wavemetrics, Large Oswego, OR) for the analysis of electrophysiological data, and NIH ImageJ/Fiji software [Schindelin et al. (2012)] for imaging data. Normally distributed data was expressed as mean ± SD and compared using one- or two-sided Student’s t-tests. Data was considered significantly different for P values smaller than 0.05. *, **, *** are shorthand for P < 0.05, 0.01 and 0.001 respectively. Absolute p-values are indicated in the figure legends.

## 3 RESULTS

Goertzen and Veh identified, in rat, two potassium-channel accessory proteins, KChip3 and Kv2.2, respectively, as cell-type specific markers for Fañanas cells (FCs) [Goertzen and Veh (2018)]. We successfully reproduced their experiment in rat (Fig. S2), yet neither our attempt to label FCs through immunocytochemistry nor immunofluorescence staining against KChip3 or Kv2.2 yielded any specific staining in mice. While the reasons for this species difference remain obscure, this finding forced us to revise our strategy and explore alternative approaches for investigating FCs.

### 3.1 FCs express AldH1L1 and are present from P5 on throughout adulthood

Despite the recent discovery of some unexpected non-glial expression [Foo and Dougherty (2013), Boesmans et al. (2014)], AldH1L1 has been one of the most consensual pan-glial marker since its initial description [Cahoy et al. (2008), Yang et al. (2011)], and it is increasingly being used over the traditional and more heterogeneously expressed glial fibrillar acidic protein (GFAP), the expression of which shows significant differences among brain regions and even cells within the same region [Kálmán (2002)]. Imaging sagittal cerebellar slices prepared from a transgenic mouse line expressing AldH1L1-GFP selectively in astrocytes [Cahoy et al. (2008)] we examined whether FCs expressed this astroglial enzyme. Indeed, in addition to Bergman glia (BGs), confocal micrographs revealed abundant GFP-positive cells with small cell bodies located in the molecular layer (ML) and exhibiting the characteristic dart-like morphology of FCs (Fig. 2A). This immunolabeling against AldH1L1 comforted observations (Fig. S3) and it confirmed AldH1L1 expression by the feathered cells of the cerebellum. Compared to alternative markers for astrocytic identification, ALDH1L1-GFP demonstrates minimal background noise within ML. This is attributed to its relatively low expression levels in the extensions of BGs. Conversely, FCs exhibit robust expression of ALDH1L1-GFP, thereby facilitating a pronounced contrast of FCs in our acquired images. Interestingly, FC counts in adult slices (an example at P27 is shown in Fig. 2A) revealed marked differences in cell density between lobules (Fig. 2B), and these differences were consistently observed at all ages (Fig.2C). We reliably detected FCs from P6 on and throughout adulthood. Pooling cell counts from all lobes, we observed no significant change in FC density with age (Fig. 2D). Despite some variability between animals, FC density was stable through adulthood, indicating that FCs remain in place and are not just displaced BGs that failed to integrate into the cerebellar circuit and are bound to disappear with age.

**FIGURE 1.**
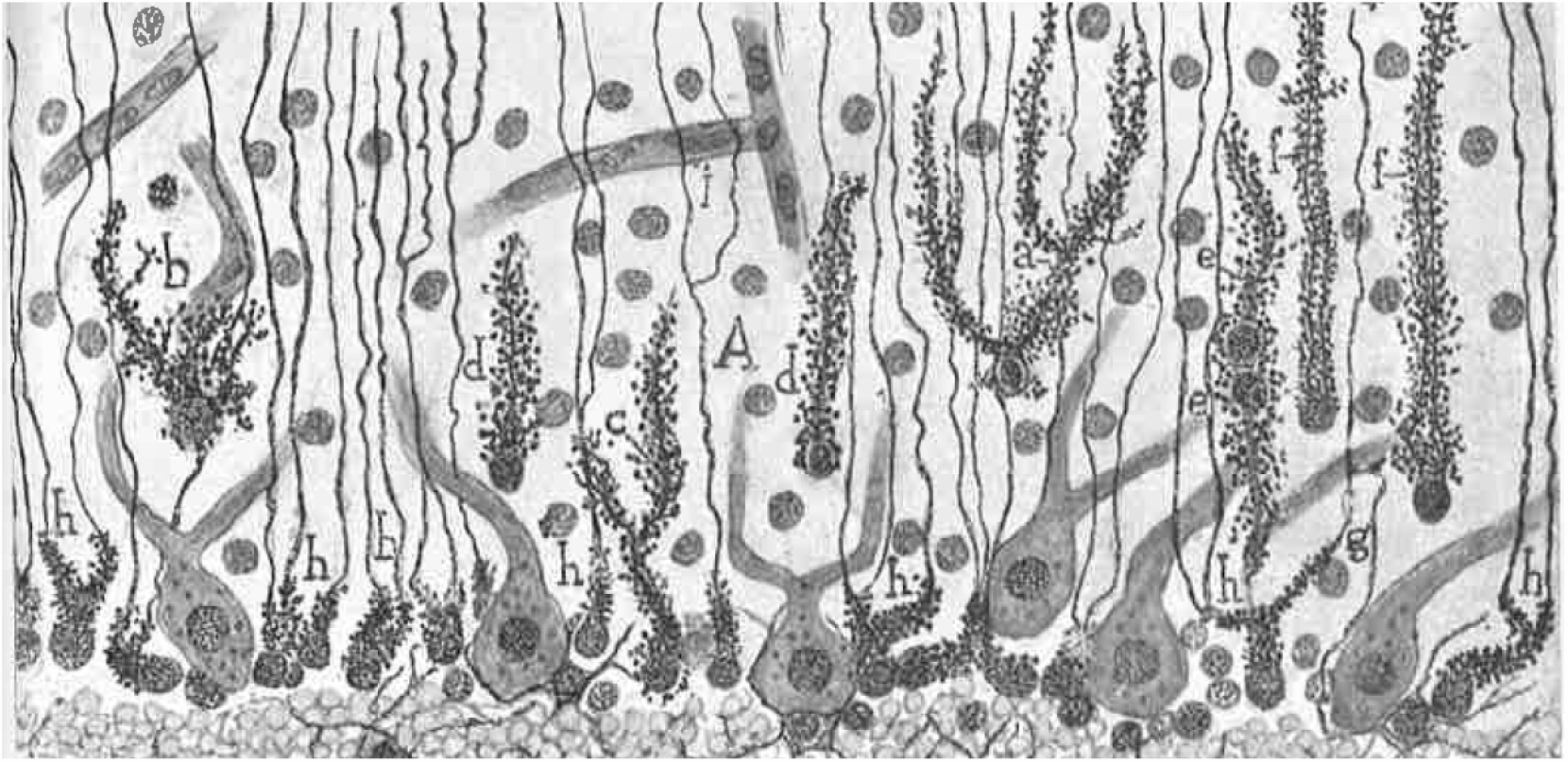
First Fañanas cells representation from histological gold-sublimation method by Ramón y Fañanas in 1916. In this drawing we can distinguish Purkinje cell, Bergmann glia (h) and Fañanas cells (d or b)

**FIGURE 2.**
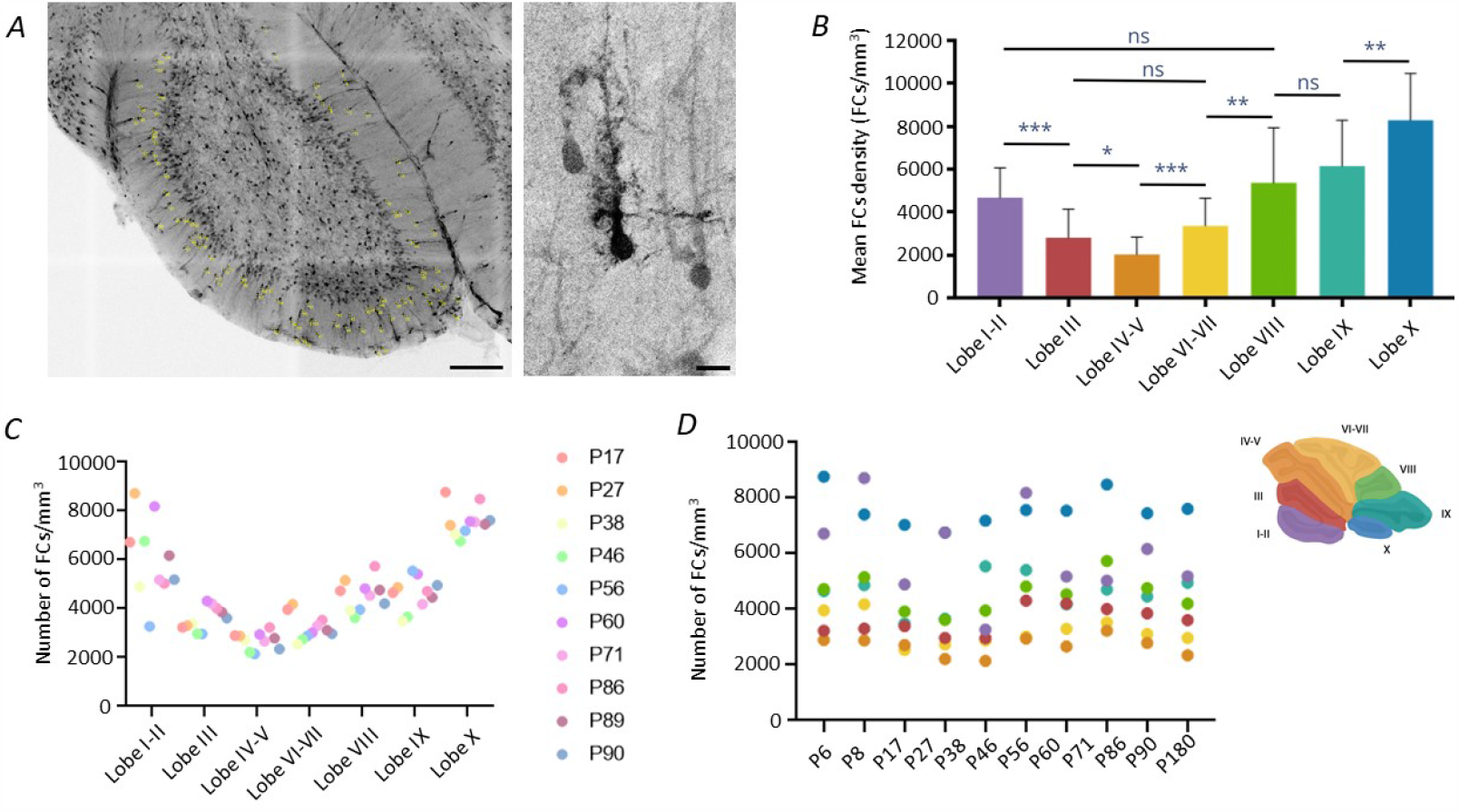
The distribution of Fañanas cells in the sagittal molecular layer varies between the different lobes. A) confocal image of a 200 μm thick cerebellar slice of an adult Aldh1-EGFP mouse (P27) with supposed FC indicated with yellow dot. Images taken at x20/0.5 are a sub-stack of 30 μm zoomed on lobe X, scale bars are 100 μm for the left panel and 10 μm for the right panel. The right panel focus on a single FC. Image taken with Zeiss x63/1.4 oil immersion lens (WD: 0.19 mm). B) the number of FCs varies between lobes, with the lowest density consistently found in lobes IV-V and the highest in lobe X. Cerebellar sub-divisions used for counting are shown in the top right (n = 11 animals and n = 2 to 3 slices per animal). C) Data for every mice counted showing that the same profile is found in all animals from P17 to P90. D) The total cell density of FCs does not vary significantly during adulthood.

### 3.2 FCs have passive membrane properties similar to those of other astrocytes

After some experience, we were able to readily identify FCs on DIC images and patch them without confusion with interneurons, even in wild-type (WT) mice, with no need for systematically using Aldh1L1 reporter mice. In a first set of experiments, to better visualize individual FCs on a dark background, we added Alexa 594 dye into the intracellular solution and we imaged slices immediately fixed after filling using confocal fluorescence microscopy. Figure 3A shows an example of such a 3-D reconstruction. FC morphology was very reminiscent of that seen in the original drawings by Ramón y Fañanas (see Fig.1), lending further support to our identification. In comparison to neighboring Bergman glia (BGs) (Fig. 3B), FCs typically had a simpler morphology with one or, at most, two stubby processes (1.3 ± 0.4, n = 12) that measured, on average, approximately half the length of the highly branched, candelabrashaped processes of Bergman glia (Fig. 3C; 60.5 ± 22.9 μm, n =12 for FCs vs. 149.5 ± 11.1 μm, n = 12 for BGs).

**FIGURE 3.**
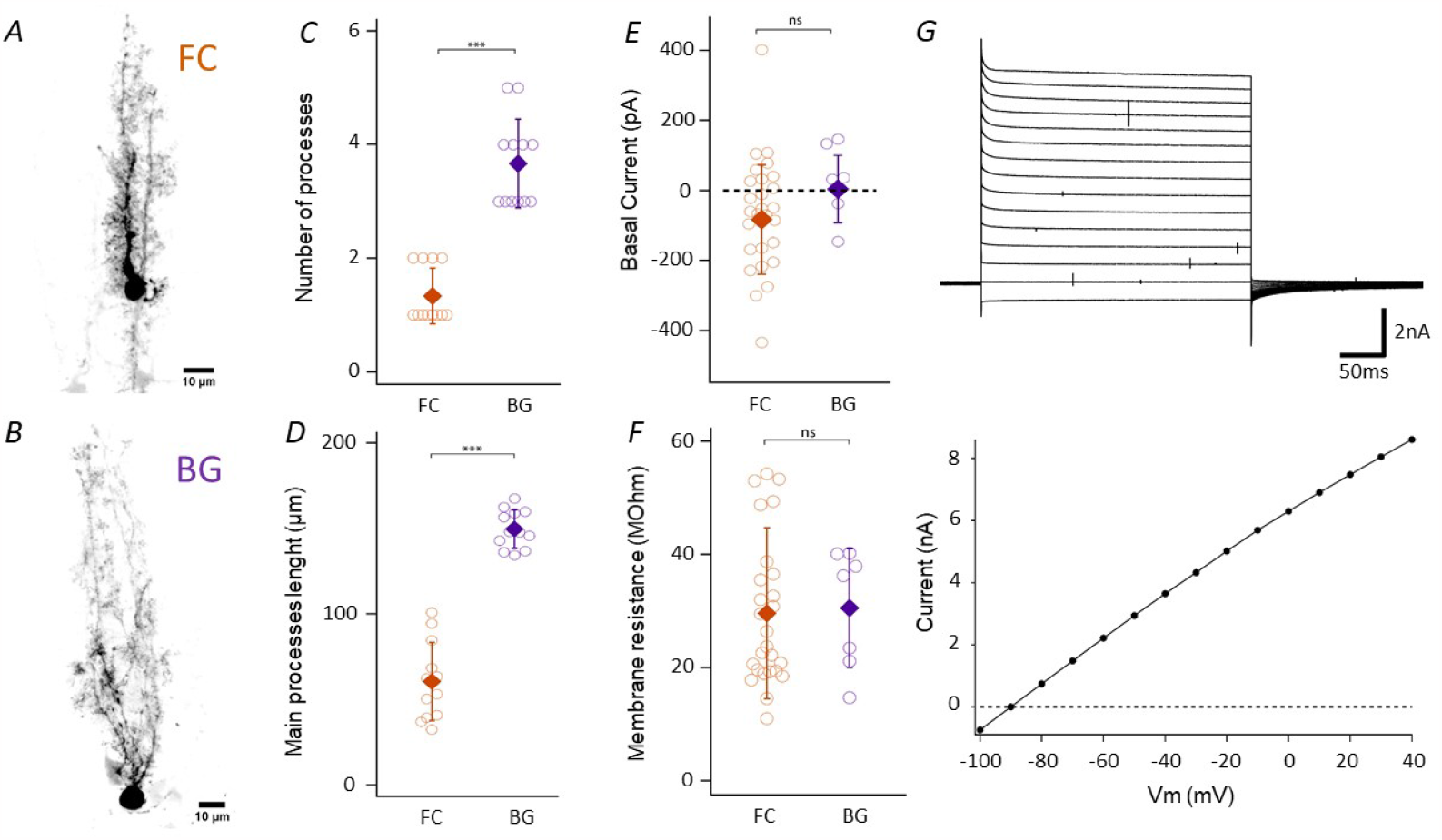
FCs are morphologically different but electrophysiologically indistinguishable from BGs. A) FC confocal construction of a FC after filling with Alexa 594. B) Same for BG. C) Comparison of the number of principal processes in FCs and BGs (n=12 cells in 7 mice). D) Maximal process length from the soma (n=12 in 7 mice). E) Currents recorded at -90 mV from FCs (n=26 in 16 mice) and BGs (n=7 in 7 mice). F) Membrane resistance calculated from the currents elicited by a -20 mV pulse applied form a holding potential of -90 mV. G) Top, whole-cell recorded currents upon voltage steps from -100 to 0 mV, 10 mV steps; bottom, the resulting IV curve indicating dominantly passive membrane conductances and current reversal near -90 mV, close to the K+ reversal potential.

Upon establishing the whole-cell configuration, FCs displayed series resistances (R_s_) of approximately 30 MΩ, and a membrane capacitance around 20pF. In the absence of stimulation, holding currents were around 400 pA at -60 mV and -66 ± 162 pA at -90 mV (Fig. 3E). We only included cells for which R_s_ did not change more than 15 MΩ during recording. I-V curves from -90 to 0 mV (Fig. 3G) revealed largely passive conductances with membrane resistance of 29.6 ± 13.2 MΩ (11 to 54.2 MΩ, n = 26 FCs vs. 30.5 ± 10.5 MΩ, n =7 BGs) (Fig. 3F). Based on these observations, we decided to keep cells at a holding potential of -90 mV, close to the potassium equilibrium potential. Results did not much vary at 34°C, suggesting the temperature independence of these passive membrane properties (Fig. S4).

Based on our electrophysiological data, we conclude that although FCs are noticeably smaller and have a more compact cytoarchitecture, they are indistinguishable from BGs in terms of membrane electrical properties. Similar to their larger counterparts, FCs’ resting membrane properties appear to be primarily influenced by potassium conductances and, notably, we found no evidence of voltage-gated channels. We note that in either case, the presence of large conductances likely limited full voltage control. One possible explanation is that FCs are coupled to other glial cells through gap junctions, as previously reported for BGs [Müller et al. (1996)]. We explore below this possibility later, in section 3.5.

### 3.3 Parallel-fiber stimulation generates biphasic, AMPA-mediated currents in Fañanas cells

The cerebellum receives its main inputs from the cerebellar cortex, spinal cord and the vestibular system, resulting in granule cell activation followed by parallel fibre excitation. To mimic this physiological excitatory input, we placed an extracellular stimulation electrode in the molecular layer (ML), while recording from a single FC, Fig. 4A. FCs responded to a stimulation train (0.11ms pulse duration, 20V, 10 pulses at 100 Hz) with a transient, biphasic inward current Fig. 4B, top, which was largely attenuated when blocking AMPA receptors with 2,3-dioxo-6-nitro7-sulfamoyl-benzo[f]quinoxaline (NBQX, 2 μM), similar to what was observed in BG, Fig. 4B, bottom. The evoked currents (37.5 ± 21.6 pA for FCs and 41.2 ± 29.2 pA for BGs, Fig. 4C) and fractional current block by NBQX (77.5% of CTR, n = 9 for FCs vs. 76.8%, n = 6 for BGs) were indistinguishable between FCs and BGs (Fig 4D).

**FIGURE 4.**
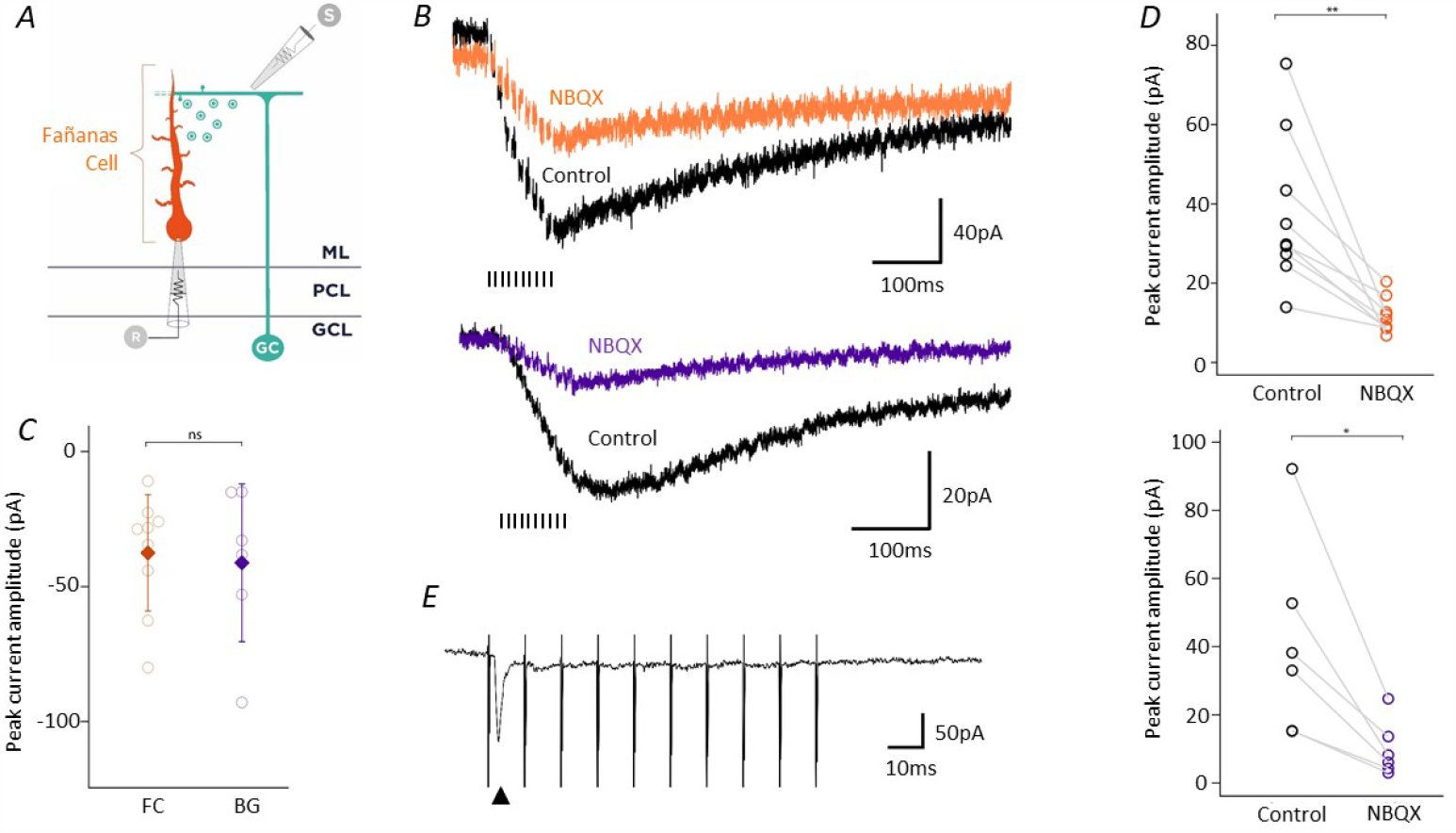
Parallel Fiber stimulation generates monophasic AMPA-mediated currents in Fañanas cells. A) Schematic representation of extracellular electrical stimulations and whole cell recordings. B) Quantification of current amplitudes at the end of the stimulation train in BGs (n=6 in 6 mice) and FCs (n=9 in 8 mice) C) FC whole-cell recording during molecular layer stimulation, before and after NBQX treatment. We applied 100 Hz stimulation at 40V. The same recording has been done in BG for comparison in the bottom panel. D) Comparison of the peak amplitude during PF train stimulation before (37.5 ± 21.6 pA) and after NBQX (8.5 ± 4.9 pA). Upper panel: FCs (n=9 in 8 mice) (p-value = 0.0011). Lower panel: Same experiment on BGs (before NBQX: 41.2 ± 29.2 pA; after NBQX: 9.6 ± 8.3 pA) (n=6 in 6 mice) (p-value = 0.0288). E) Fast current (1.1 ms rising time) during PF stimulation (arrowhead), observed in 2 cells out of 9.

As expected, very similar results were obtained when placing the stimulation electrode in the granule cell layer (GCL) instead of the ML (Fig. S5). Thus, similar to what has been described for long in BGs [Clark and Barbour (1997)], FCs express functional AMPA receptors and respond to extracellular glutamate. We noted, for both FCs and BGs the occurence of occasional fast-amplitude inward currents superimposed on the slower current envelope as if FCs - again like BGs - [Clark and Barbour (1997)] detected synaptic inputs in addition to ambient glutamate (Figs. 4E, arrowhead).

In response to repetitive PF stimulation at 100 Hz, FCs showed a marked attenuation of the current amplitude. Therefore we decided to investigate how the precise release of glutamate affected AMPA-mediated currents.

### 3.4 FCs detect fast glutamatergic inputs on the millisecond-time scale

Parallel-fiber (PF) stimulation produces a complex spatiotemporal glutamate concentration profile, which makes the evoked current difficult to interpret. To better control glutamate release, we therefore turned to near-UV laser photolysis of caged MNI glutamate [Trigo et al. (2009)], Fig. 5A. Glutamate uncaging allows for a localized glutamate release with a fast, step-like kinetics due to the decoupling glutamate diffusion and release. We did not observe deleterious side effects from photolysis by-products. The calibrated laser power and the large reservoir of caged MNI glutamate in the extracellular space produces high reproducibility (Fig. 5B). Indeed, in sagittal cerebellar slices bathed in 0.5-1 mM MNI glutamate, laser photolysis induced in FCs rapid currents that mimicked the fast inward currents observed previously with extracellular stimulation, with a similar depression of peak amplitude upon repetitive photolysis than that earlier observed in response to pulse trains (Fig. 5C).

**FIGURE 5.**
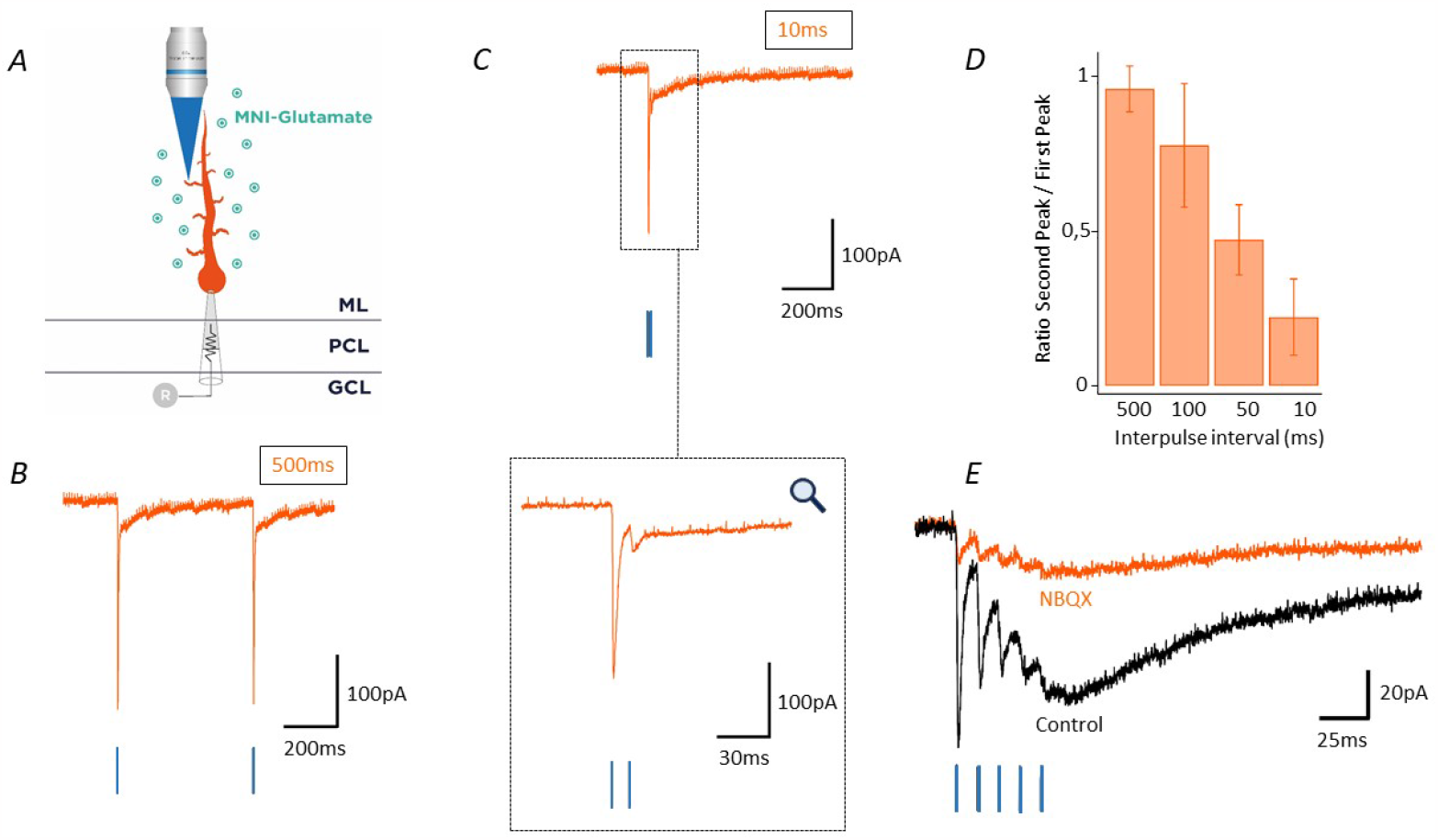
Glutamate uncaging on FCs triggers a biphasic AMPA-mediated current. A) Schematic representation of the photolysis configuration of MNI-glutamate on FCs. B) A 405 nm laser twin pulse (500 ms interval) on a main FC process triggers fast and strong peak currents. C) The same twin laser pulse but with a 10 ms interval leads to a strong amplitude reduction of the response elicited by the second pulse. The bottom panel shows the same data but with an expanded time scale to highlight desensitization of the second peak. D) Desensitization quantification during different light pulse intervals between 500 to 10ms (n = 5 in 5 mice) E) Currents elicited by a 5 near-UV pulse train at 100 Hz before and after NBQX treatment observed in two cells. In this experiment, the laser power was lowered to reach a small current, similar to that elicited by electrical train. NBQX decreases by 85.4% of the fast current and 75.3% the slow current.

The observed paired-pulse depression (PPD) depended on the inter-pulse interval, with a full recovery of the current peak amplitude only for pulses more than 0.5 s apart (Fig. 5D). On the contrary, shorter pulse intervals resulted in a rapid onset of AMPA-receptor desensitization. Notably, an interval below 10 ms effectively blocked the triggering of the second pulse. When examining inter-pulse intervals ranging from 500 ms down to 10 ms, we observed a linear decrement in the efficacy of the second pulse. This suggests a systematic attenuation of the response, possibly indicating a temporal refractory period or a mechanism of synaptic adaptation. Hence, with an appropriately chosen set of glutamate release and inter-pulse interval, we successfully replicated with near UV-laser-uncaging the earlier demonstrated extracellular train stimulation (Fig. 5E). As upon extracellular stimulation we observed a slowly rising current in response to glutamate. However, when we applied glutamate directly onto the cell, we systematically observed an additional fast current that exhibited a fast desensitization, occasionally observed upon extracellular stimulation, (cf Fig. 4D). The AMPA receptor antagonist NBQX completely blocked this fast current, underscoring its dependence on AMPA receptor activity. In contrast, the slowly-rising current displayed only a partial suppression under the influence of NBQX. This discrepancy in NBQX sensitivity between the two current types suggests distinct underlying mechanisms or the involvement of different receptor subtypes.

Next, by systematically displacing the laser spot along the FC process, we mapped glutamate-evoked currents in a spatially resolved manner (Fig. 6A). We observed the largest current amplitudes in the proximal process (Fig. 6B), even after normalizing the measured currents to those recorded at the somatic level in the same cell (Fig. 6C). However, considering that the size of the UV spot is smaller than both the cell body (position “1”) and the proximal process (“2”), but larger than the size of the fine protrusions emanating from the secondary processes that are not resolved in the epifluorescence image, it is reasonable to presume that local current densities (and, by extrapolation, AMPA receptor densities) were highest in the distal processes. This interpretation aligns with the observed 3-D architecture. Confocal micrographs that offer a higher spatial resolution than the epifluorescence images acquired during photolysis and revealed a conspicious proximity of FC processes with PC dendrites (not shown). Thus, albeit indirect, our electrophysiological, uncaging and morphological data together suggest that FCs are capable of monitoring synaptic activity. Further ultrastructural studies are necessary to elucidate the precise nature of these neuron-glia contact sites and functionnal studies must clarify the involved signaling pathways.

**FIGURE 6.**
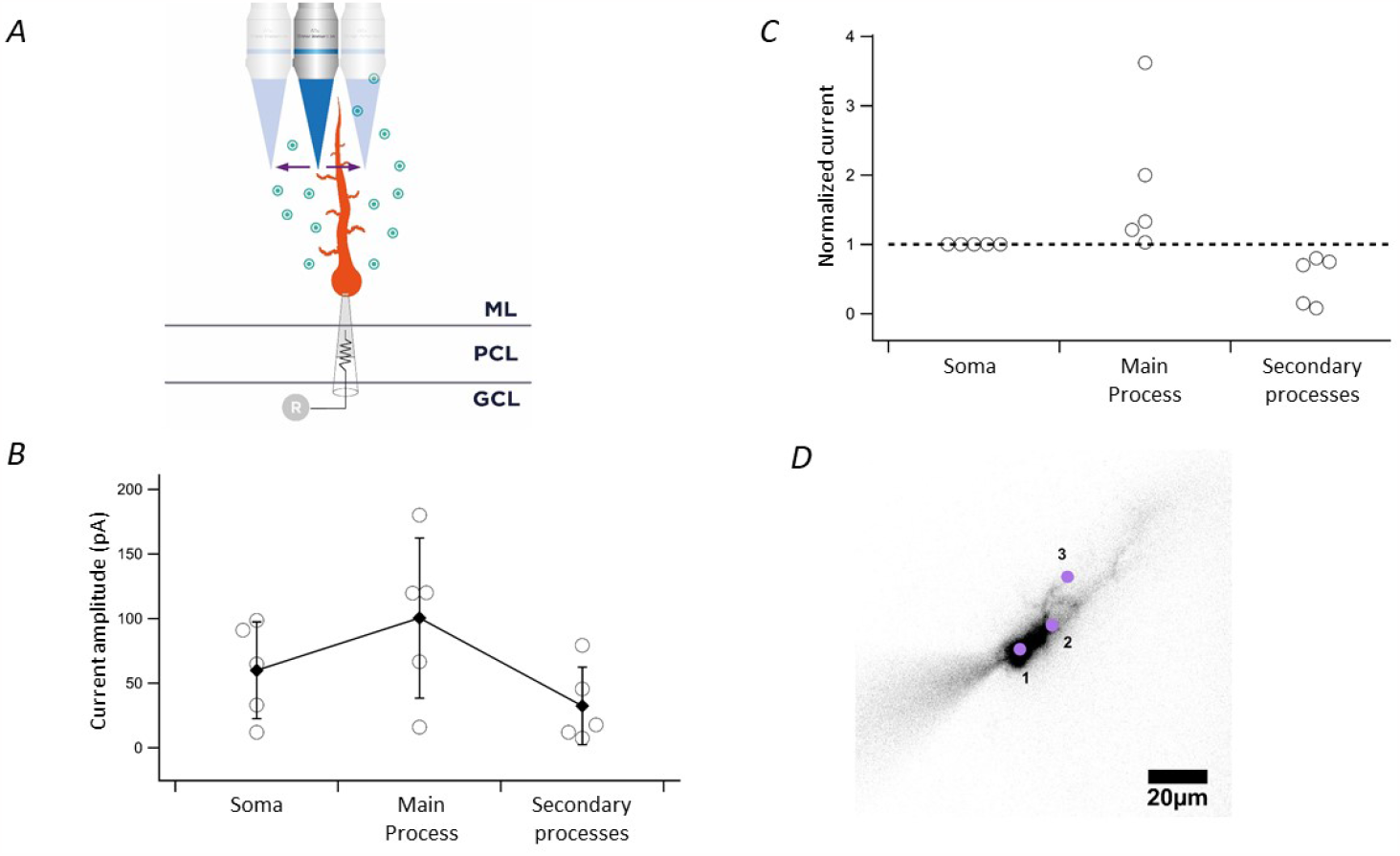
Glutamate receptors are ubiquitously expressed on FCs with higher density on process membrane. A) Schematic representation of the photolysis configuration for MNI-glutamate release on FCs. B) Current amplitude during targeted photo-release of glutamate on 3 cell regions: the round soma, main thick process and small processes (n=5 in 5 mice). C) Data from A normalized to the soma (n=5 in 5 mice). D) Epifluorescence imaging of FC after 10 minutes filling with Alexa-594. Typical regions for stimulation are indicated with purple dots.

### 3.5 FCs integrate into the glial syntitium

We earlier found that FCs and BGs displayed similar passive electrical properties, despite their distinct morphologies (Fig. 2). To better understand this at first sight puzzling observation, we next evaluated the extent of gap junction coupling of BGs and FCs by a combination of whole-cell patch clamp recordings and dye filling/spreading experiments in the presence of drugs that reduced or increased, respectively, the amount of gap-junction coupling.

First, similar to the experiment shown in Fig.2, we recorded current in response to PF stimulation in either FCs or BGs. Currents recorded in either cell type were sensitive to Carbenoxolone (CBX), a well-known gap junction inhibitor. 150 μM CBX reduced the measured peak current amplitude by 50% (Fig. 7A and B), and this consistently across all tested cells, with the mean current amplitude decreasing from 209.4 ± 16.7 pA to 119.5 ± 43.7 pA (Fig. 7C). Moreover, CBX decreased the membrane conductance (see Fig. 7D for FC data), strongly supporting the hypothesis that both FCs and BGs are interconnected through gap junctions. To further substantiate this observation, we introduced 0.8 μM spermine into our internal solution to open gap junctions. As expected, we observed that Alexa-594 dye included in the intracellular solution spread to a large number of neighboring cells, and - based on the immediate fixation and 3-D reconstruction after a 20-min filling episode - we recognised both FCs and BGs among the filled cells (Fig. 7E). Together, this data provides clear evidence for a robust homo- and heterotypic connectivity between FCs and BGs through gap junctions, underscoring the role of inter-cellular coupling in their common electrical properties. Thus FCs functionally integrate into the glial syntitium.

**FIGURE 7.**
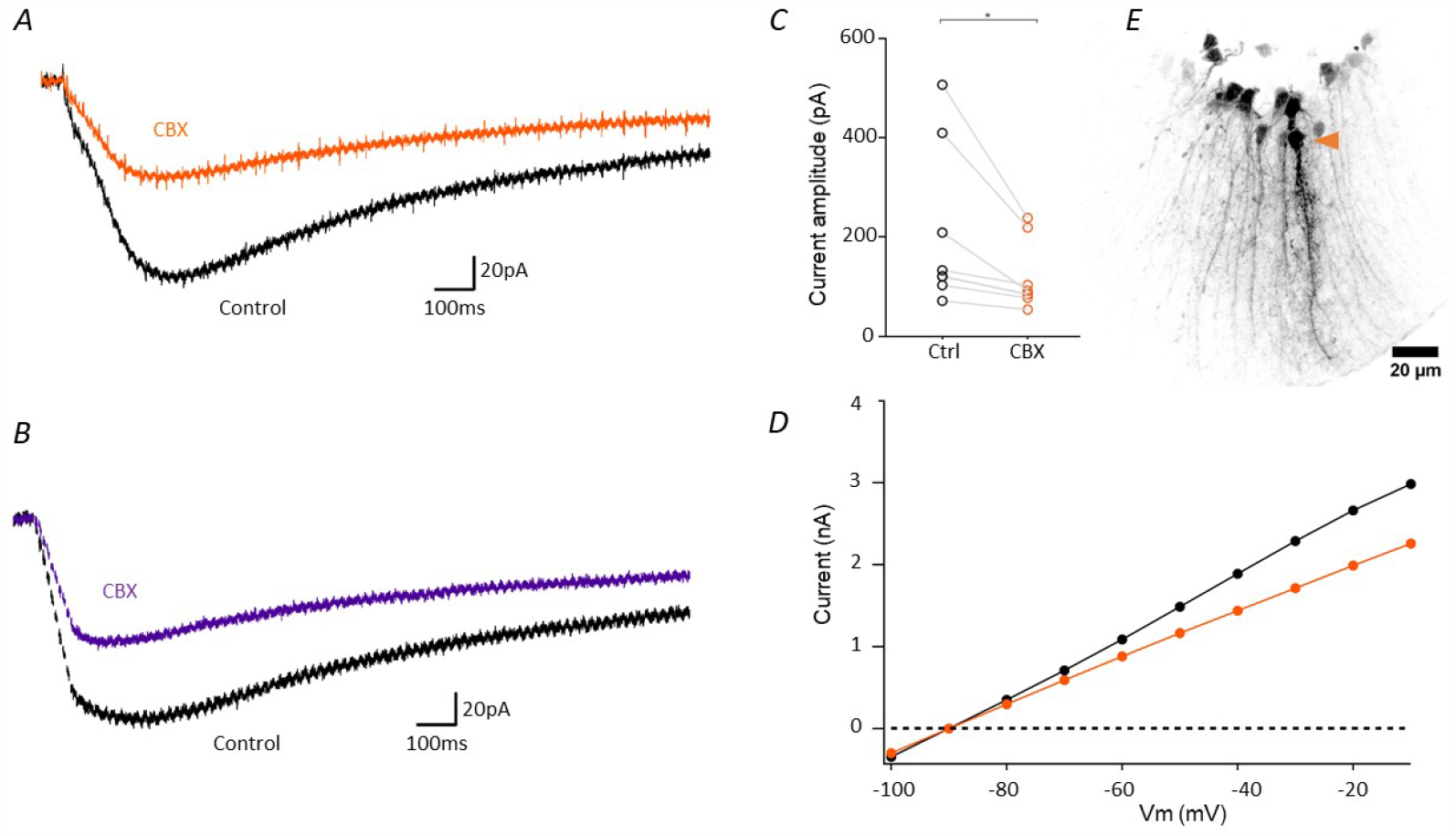
Gap junction coupling in FCs. A) FC recording during PF stimulation before and after 150 μM CBX. B) Same experiment on BG for comparison. C) Quantification of CBX effect on FC during PF stimulations. D) IV before and after CBX from -100 to 0 mV. E) FC filling with 0.8 mM Spermin in internal solution leads to opening of gap junctions. The FC initially recorded from is indicated by the orange arrowhead.

In a final experiment we took advantage of the high spatial resolution of the uncaging technique and tried to combine it with the recording of a pair of FCs coupled through a GAP junction. In order to do this experiment we included spermine in both internal solution, as indicated above. During one experiment, we found two closeby FCs and realized a double recording, indeed both cells were electrically coupled and the results of this experiment are presented in Figure S6. Panel A shows the results of depolarizing the magenta FC and recording B the opposite. A current can be recorded in the respectively non-stimulated cell confirmed their electrical coupling. Panel C and D show a similar experiment, but now, instead of applying voltage pulses, glutamate was photoreleased in either one FC (the black one in C or the magental one in D). The current recorded in the non-tested, electrically coupled cell was not due to a direct effect of the photolysed glutamate because it disappeared when the laser spot was moved away from the stimulated cell (data not shown). Some very interesting conclusions can be drawn from this experiment. First, as suspected from own earlier results, the glutamate released onto one FC induces a current in this cell, but also a significant GAP-junction dependent current in the neighboring FCs (and probably in all the other cells that are part of that syncytium). Second, as expected from the known properties of GAP junctions, to act as a low-pass filter, the fast response is much more attenuated than the slow one. In the example shown in Fig. S6D, the fast response had an amplitude of 544 pA in the stimulated cell, where glutamate was photolyzed, and 170 pA in the coupled one (a 70% attenuation) while the slow response passed from 56 pA in the stimulated cell to 53 in the post-synaptic one (almost no attenuation). This can also be seen when we used a train of near-UV laser pulses to photorelease glutamate on the stimulated cell (Fig. S6E, left and right). These data supports the view that FCs are part of a glial syncytium, and the glutamatergic currents in FCs are a complex mixture of currents originating in the membrane of the FC and currents originating in other, GAP-junction coupled cells.

## 4 DISCUSSION

### 4.1 Fañanas cells, the long neglected astrocytes of the cerebellum

Fañanas cells, are a type of cerebellar astrocyte, primarily located in the granular and molecular layers, with their protrusions extending into the lower part of the molecular layer [Goertzen and Veh (2018)]. FCs have been considered to be closely related to, and sometimes even identified as Golgi epithelial cells [Chan-Palay and Palay (1972)]. Some researchers have classified FCs as “specialized astrocytes”, while others regarded them as just a shorter variant of BGs [Reichenbach and Bringmann (2020)]. FCs owe their name to Jorge Ramón y Cajal Fañanas, the son of Santiago Ramón y Cajal, who was the first to describe this type of glial cell in 1916 [Fañanas (1916)]. Although discovered over 100 years ago, FCs have received a very limited attention in scientific literature. A comprehensive literature search yielded only 49 relevant mentions (as of June 2023), many among them being reviews, citations, or perspective articles, and only a few represent original research articles. Detailed morphological and functional data on FCs has been missing.

### 4.2 Possible functions of Fañanas cells

The role of FCs in the cerebellar circuitry is unknown. Our results seem to indicate a role for FCs other than just a displaced BGs: they are and remain present in the mouse brain at all ages and they are functionally integrated into the circuit, and this by at least two mechanisms. First, and perhaps with little surprise, they fulfil classical astrocyte housekeeping activities, as suggested by their dominantly potassium channel conductances, (Fig.2) their response to ambient glutamate with a slow inward current (Fig. 4) and their functional integration into the astroglial syncytium via gap junctions (Fig.7). At the same time, FC AMPA receptors read out fast (ms) glutamate release with high fidelity (Fig. 4) and thus FCs are potentially endowed to detect local synaptic activity. The morphological proximity of the FC process and PC dendrites further points into this direction, and we will investigate this topic further in a second paper (Singer*, Vinel* et al., *in preparation*).

Like BGs, FCs functionally express AMPA receptors blocked by NBQX (Fig. 4) and they respond to repetitive PF stimulation with a rapidly desensitising, biphasic current (Fig.5). The localised fast glutamate release produced by uncaging MNI-gultamate mimicked this behaviour and reproduced notably the fast current spikes that we occasionally observed super-imposed on the slow inward current. Together with our observations of intracellular coupling (Fig. 7 and S6); the slow current recorded may be an indirect current coming from another, electrically coupled cell. Thus, taken together, our electrophysiological and uncaging data are compatible with the hypothesis of Monteiro who speculated that FCs might be a specialized satellite cell to PCs [Monteiro (1983)], although that article classified them as a type of oligodendrocyte. An additional argument against FCs being just a developmental ‘leftover’ during cerebellar circuit formation is the presence of FCs from early postnatal age and throughout adulthood. Our systematic analysis of FC number in AldH1L1-GFP reporter mice revealed marked inter-lobe differences, but a fairly constant cell density independent of age for mice after P10 (Fig.2).

### 4.3 FCs express functional AMPA receptors and are part of the glial syncytium

NBQX application reduced in FCs the amplitude of PF-stimulation evoked currents by 70-80%, similar to what was observed in neighboring BGs (fig.4). Future experiments must clarify if FCs express the same two AMPA-receptor subunits, GluA1 and GluA4, that dominate in BGs where they mediate Ca^2+^ influx [Lim et al. (2021); Saab et al. (2012); Porter and Mccarthy (1997)]. This is particularly interesting as in BGs, the viral overexpression of GluA2 renders AMPARs Ca^2+^-impermeable, impairs the glial ensheathment of PC synapses by BG processes and slows reduced glutamate clearance [Iino et al. (2001)], an observation still to confirm for FCs. In addition to their necessarily local action at the level of neighboring PC dendrites (given the small size of FCs, Fig.2), we demonstrated FCs are both electrically and diffusional coupled to neighboring glia (fig. 7E), and this in a homotypic (FC-FC) and heterotypic manner (FC-BG). Thus, similar to BGs and astrocytes in other brain regions, they are part of the large syncytium of glia coupled by gap junctions. The degree of FC coupling could be modulated in a bidirectional manner by means of spermine and CBX application, respectively, (fig. 7), and it will be interesting to study, if this coupling can be influenced through the activation of Ca^2+^-permeable AMPA receptors. Ca^2+^ regulates gap-junctional coupling in BGs in a bell-shaped manner, which has as an effect that moderate Ca^2+^ loads following stimulation are efficiently distributed among many cells, but that cellular coupling is abolished, and the network protected from pathologically high Ca^2+^ elevations [Müller and Kettenmann (1995)]. For FCs that are part of the glial network, we expect a similar regulatory loop. Future combined patch-clamp recordings and 2-photon Ca^2+^ imaging using genetically encoded Ca^2+^ indicators (GECIs) must clarify this hypothesis.

### 4.4 FCs are present throughout adulthood at fairly constant number

Contrary to common belief, FCs are not rare. Taking the number of Aldh1L1-positive cell bodies in the ML as a proxy, we found an average cell density of > 4,000 FCs/mm^3^, varying from around 2000 FCs/mm^3^ in lobe III and 2000 FC/mm^3^in lobes IV-V to about 8,000 FCs/mm^3^ in higher-density regions (anterior, posterior and nodular). Our finding that FC cell densities vary among lobes opens interesting perspectives for future research. It will be important to correlate these “hot spots” of FCs with other structural markers that report on sub-cerebellar compartments, like zebrin. We speculate that FCs might be in command of local glial networks that orchestrate activity of neighboring groups of BGs and PCs. At the cellular level, FCs were if not stereotyped, but fairly similar in shape. Confocal micrographs of dye-filled FCs show a compact stubby cell. Located near PC soma, FCs occur in forms with one or rarely two “feathers” of cytoplasmatic extensions that are studded with small, rounded sprouts. The protrusions seem different from and systematically less complex and shorter than those of BGs and they often, but not always run parallel to BG fibres. Generally, FC processes measure about half of those of BGs and they are not part of the glial limitans. Interestingly, there are anecdotic reports of pathological changes in FC shape and number in two prion diseases. Both CreutzfeldtJacob disease (CJD) [Lafarga et al. (1993)] and kuru [Liberski et al. (2012)] seem to go along with a conspicuous proliferation of FCs. However, a rigorous distinction between different types of cerebellar glia was not made in these studies so that either observation could be the sign of a more diffuse astrogliosis, rather than an effect selective for FCs, but they certainly merit a reinvestigation in view of the new data. Future studies must clarify if FC shape is plastic and can be modified in response to stimulation, specific behaviours or neuropathological insults.

## 5 CONCLUSION

Our experiments revealed a significant heterogeneity in the density of FCs among different lobes of the cerebellum, suggesting their potential involvement in lobe-specific functions. Independent of their precise locus, the density of FCs is stable, indicating their continuous presence and proper integration within the astrocytic - and broader cellular - network of the cerebellum. We can, at this point, refute the hypothesis that FCs are just displaced BGs with migration defects. At the multi-cellular level, our experiments modulating gap-junction coupling demonstrates an extensive and regulated homo- and heterotypic coupling among FCs and BGs. This, along with the presence of Cx43 in FCs (Singer, Vinel, et al, in preparation), suggests the integration of FCs in the glial syncytium, thereby emphasizing the importance of considering these long-time neglected cells, both at the single-cell and network level. Notably, the methods we employed for clearing and visualizing FCs - above all the Aldh1l1-GFP transgenic reporter mouse line - have proven valuable and effective tools for future studies of FC function.

## Abbreviations

FC: Fañanas cell
BG: Bergmann glia
AMPA: alpha-amino-3-hydroxy-5-methyl-4-isoxazolepropionic acid
NBQX: 2,3-dioxo-6-nitro-7-sulfamoyl-benzo[f]quinoxaline
STED: STimulated-Emission-Depletion
GFP: Green Fluorescent Protein
CBX: Carbenoxolone
PFA: Paraformaldehyde
GFAP: Glial Fibrillary Acidic Protein
CCD: Charge-Coupled Device
ML: Molecular Layer
GCL: Granule cell layer
PC: Purkinje cell
MNI: 4-methoxy-7-nitroindolinyl
ALDH1L1: Aldehyde DeHydrogenase 1 family member L1

## Acknowledgements

We wish to thank members of the BioMedTech Facilities service unit (INSERM US36, CNRS UAR2009) for help and support, notably the personnel of the animal house, the imaging platform (Fabrice Licata) and the fine mechanics workshop (Thierry Bastien), Boris Lamotted’Incamps for critical reading early version of the manuscript. Cendra Agulhon for providing a breeding pair of the ALDH1L1-GFP reporter mouse line.

## Supporting Information

This article includes 6 supplementary figures.

