## Supplemental data for "A first morphological and electrophysiological characterization of Fañanas cells of the mouse cerebellum"

### Supplementary figures

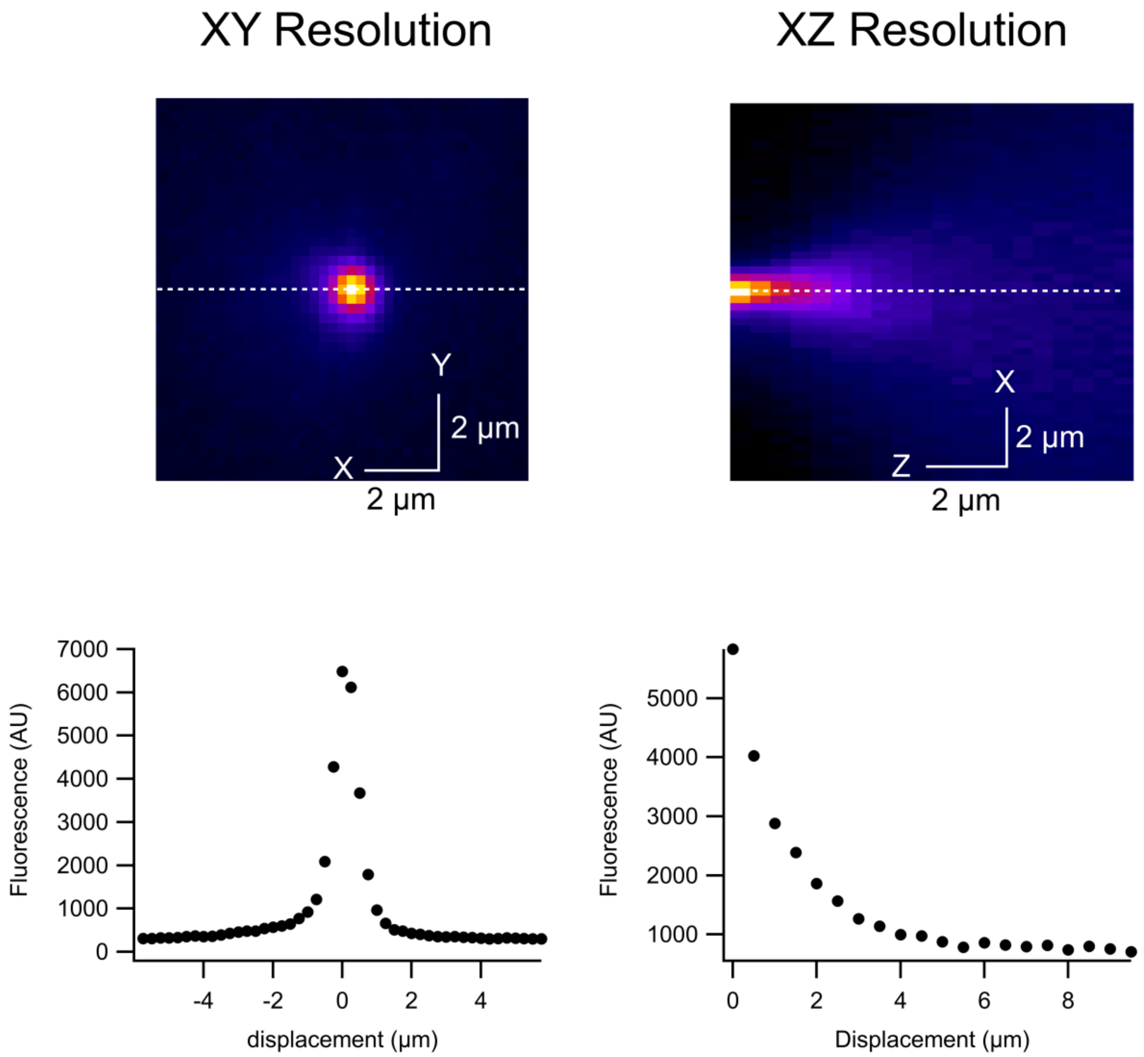


**Figure S1 : Comprehensive Characterization of the Point Spread Function (PSF).** Sub-panel A displays a microscopic image in the X-Y plane, highlighting the spatial distribution of the laser's intensity ; the color scale ranges from black (lowest intensity) to white (highest intensity). Sub-panel B shows a similar intensity-mapped image but in the X-Z plane, providing a cross-sectional view of the laser beam's focus. Sub-panel C is a graph depicting the fluorescence intensity across the X-Y plane, with the X-axis representing displacement from the center of the laser spot in micrometers and the Y-axis showing fluorescence intensity in arbitrary units (AU) ; it illustrates how the intensity diminishes as one moves radially away from the laser spot's center. Sub-panel D serves as the axial counterpart to Sub-panel C, presenting a graph of the laser's intensity as it penetrates deeper into the sample, thereby offering insights into the system's axial resolution. Together, these panels provide a multi-dimensional view of the laser's PSF, crucial for understanding its functional impact on glutamate photolysis.

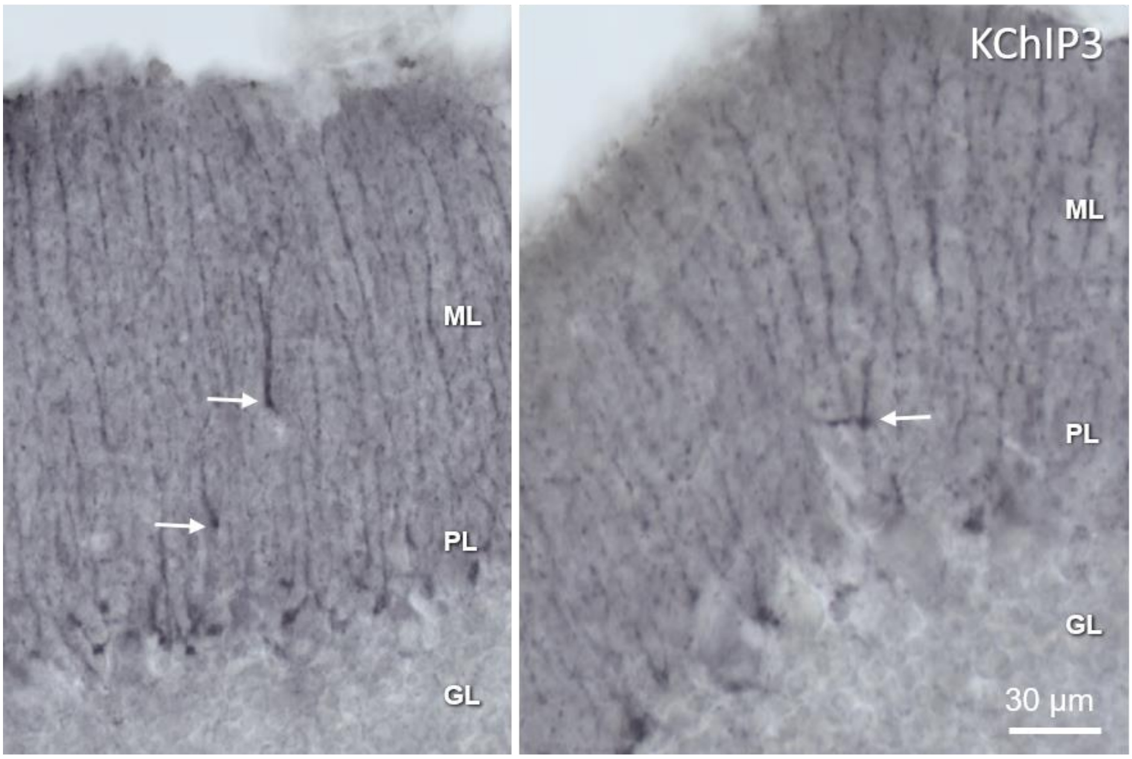


**Figure S2 : Kchip3 immunolabeling in rat.** Reproduced from Goertzen and Veh (2018)

| Lobes | Nb of cells genetic | Nb of cells immuno |
|---|---|---|
| Lobes I-II | 18 | 13 |
| Lobes III | 23 | 5 |
| Lobes IV-V | 36 | 28 |
| Lobes VI-VII | 46 | 33 |
| Lobes VIII | 53 | 19 |
| Lobe IX | 110 | 58 |
| Lobe X | 41 | 30 |

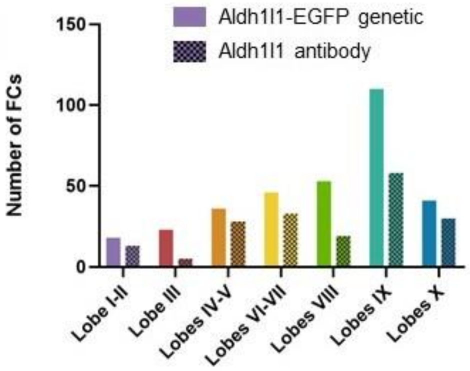


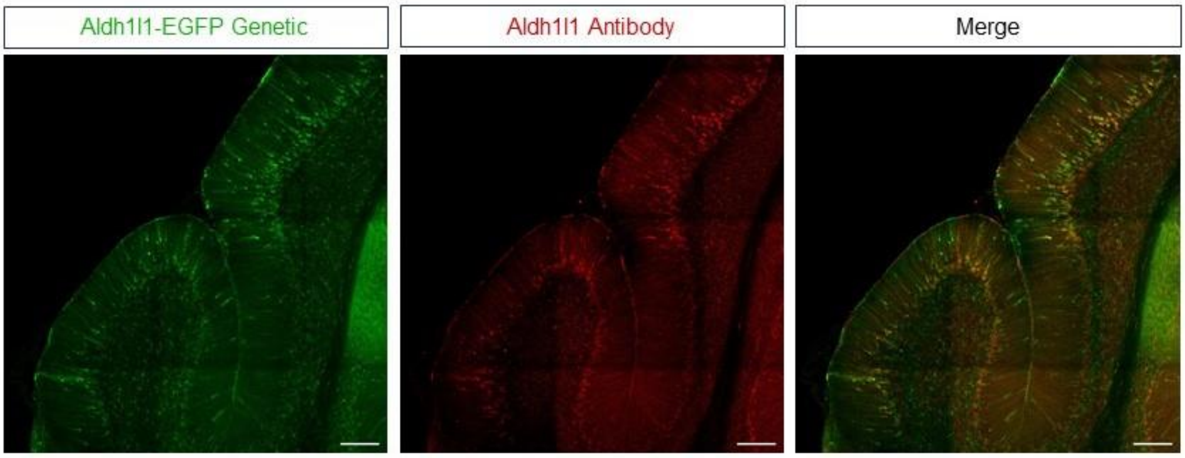


**Figure S3 : ALDH1L1-GFP mouse model validation.** Table summarizes FC counting with Aldh1l1-GFP genetically expressed compared to immunostaining against Aldh1l1. Every cells stained with antibodies expressed also Aldh1l1-GFP but the antibody did not stained all of the Aldh1l1-GFP positive cells. Right panel shows the distribution throughout lobes. The bottom panel illustrate the comparison between the 2 methods of staining observed via confocal microscope with x20/0.8 Air. Scale bar = 100 µm. For ALDH1L1 immunolabeling and confocal microscopy, we used standard techniques as stated in the main text. Used primary and secondary antibodies were XXX (dilution) and XXX (dilution), respectively.

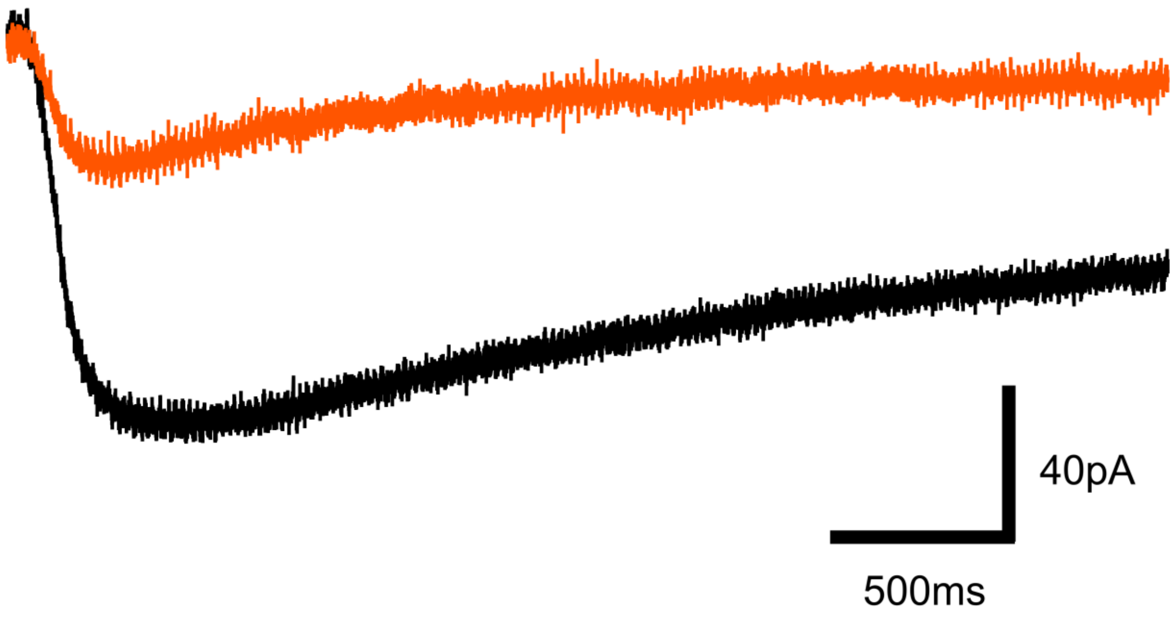


**Figure S4 : NBQX effect on FC inward current during PF stimulation.** This experiment has been done at 34°C.

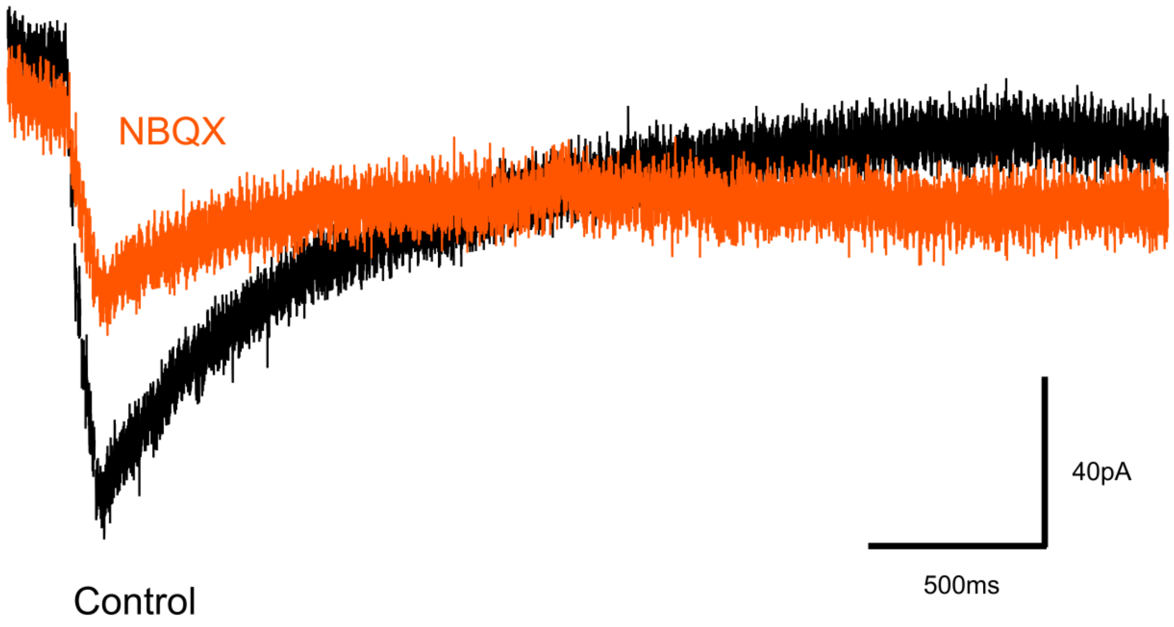


**Figure S5 :** Similar experiment to figure 4 with granular cell layer stimulation

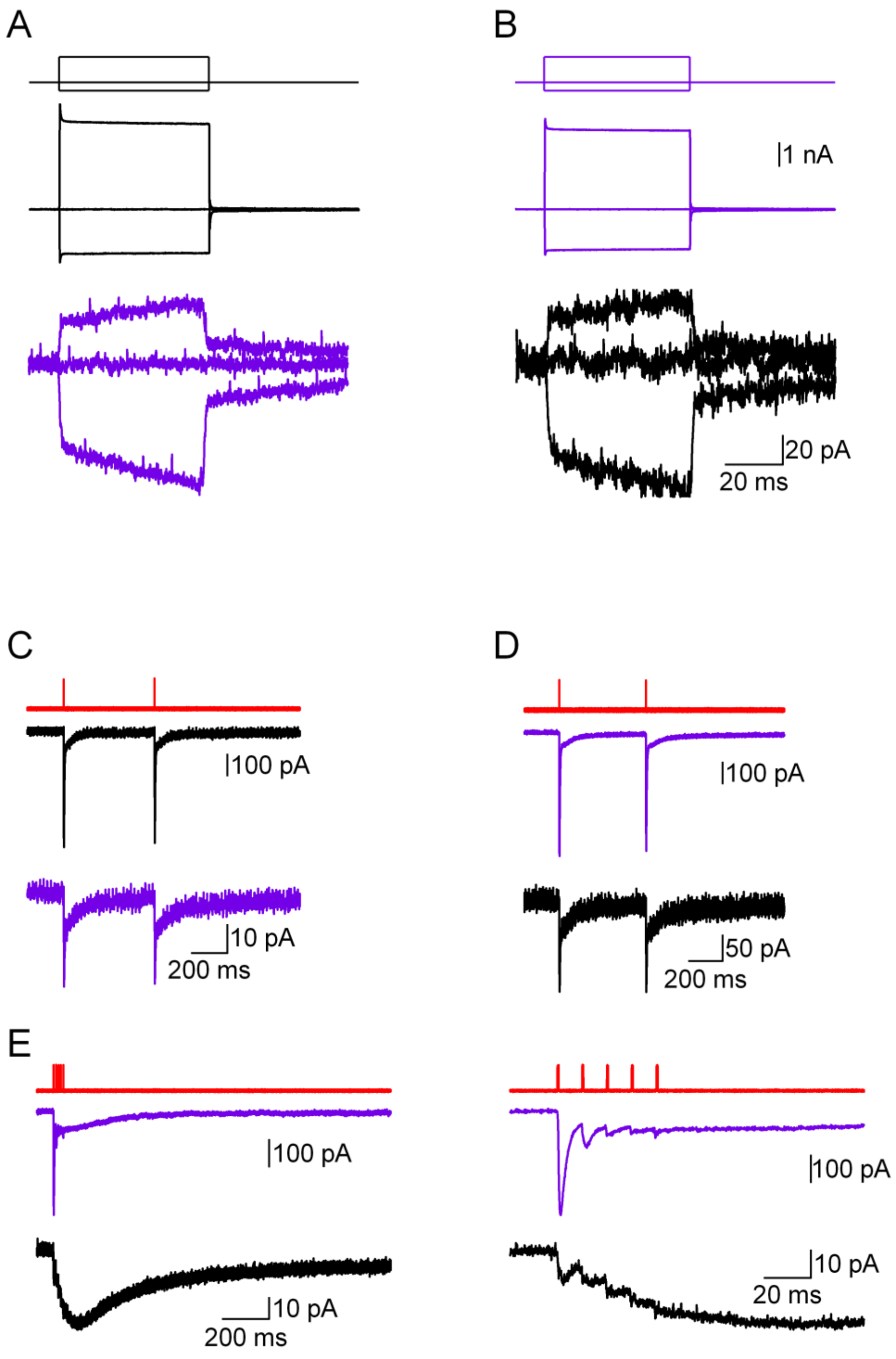


**Figure S6 : A pair of electrically coupled FCs was recorded.** A and B show the results of an experiment where the membrane currents were recorded simultaneously in both cells as a result of a series of voltage steps that were applied to only one cell (the black FC in A and the magenta FC in B. C and D show the results of an experiment were glutamate was photolysed in either the black (C) or the magenta (D) FC, while the membrane current recorded from both cells. E. Glutamate was photolysed on the black cell with a train of laser pulses. The right graph shows the same experiment with an expanded time axis. In C to D, red traces correspond to the laser pulses.
